# DOCK7 regulates WNT signaling and NMJ clustering in normal and DMD muscle

**DOI:** 10.64898/2025.12.23.696254

**Authors:** Katherine G. English, Shelby N. Rorrer, Muthukumar Karuppasamy, Kathleen A. Becker, David K. Crossman, Michael A. Lopez, Anna E. Thalacker-Mercer, Matthew S. Alexander

## Abstract

Dedicator of cytokinesis (DOCK) proteins have diverse and critical roles in myoblast fusion, glucose metabolism, skeletal muscle regeneration, and neuronal polarity. DOCK7 is part of this 11-member family of atypical guanine exchange factors. While human *DOCK7* pathogenic variants are rare, biallelic variants are known to result in hypotonia and ataxia in children. More recently, *DOCK* variants have been shown to impact disease pathogenesis as secondary modifiers. We used tissue-specific *Dock7* conditional knockout mice to evaluate the roles of DOCK7 in the skeletal muscles and motor neurons. Both *Dock7* knockout models developed significant nerve and muscle function deficits; however, the *Dock7* motor neuron knockout mice showed more severe locomotor pathologies. RNA sequencing of the *Dock7* knockout skeletal muscles revealed dysregulation of WNT signaling factors, and skeletal muscle repair and regeneration pathways. Molecular analysis revealed impaired DOCK7-mediated RAC1 activation that impacted acetylcholine receptor (AChR) clustering in both models. Interestingly, haploinsufficiency of *Dock7* in *mdx* mice improved overall dystrophic muscle pathologies. We propose that DOCK7-RAC1 interactions result in activation of the WNT signaling pathway via the LRP4 receptor in the skeletal muscle and facilitate post-synaptic AChR clustering.

**Graphical Abstract:** DOCK7 is a regulator of normal and DMD muscle via the RAC1/WNT pathway.

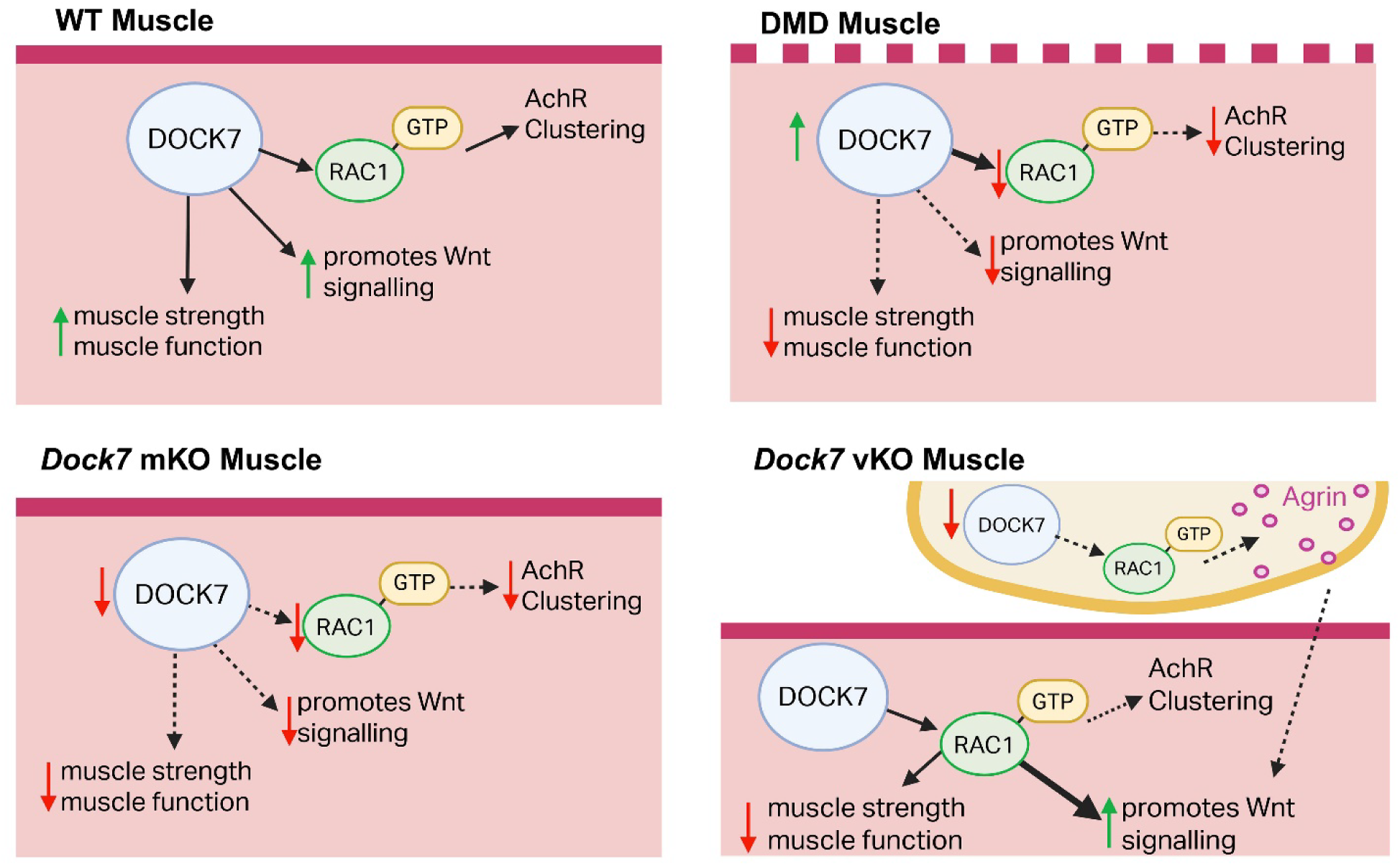

## Introduction

Dedicator of cytokinesis (DOCK) proteins are comprised of an 11-member family of atypical guanine exchange factors (GEFs) that are highly conserved in vertebrates and have orthologs in *C. elegans*. There are four subclasses of mammalian DOCK proteins (A-D) divided based on domain homology[1]. All DOCK proteins contain the DOCK Homology Region 1 (DHR1) and DOCK Homology Region 2 (DHR2) domains. As GEFs, DOCK proteins activate Rho-GTPases, such as RAC1, CDC42, or RHOA, by swapping a GDP molecule for a GTP molecule. Typically, this is accomplished by a Dbl Homology (DH) and a Pleckstrin Homology domain (PHD), and together these domains bind the inactive Rho-GTPase, catalyze the dissociation of GDP, and allow for the rapid replacement of GTP due to the ratio of intracellular GTP/GDP[2]. DOCK proteins utilize the DHR2 domain to accomplish GDP dissociation[3]. Together with GTPase-activating proteins (GAPs) and RHO guanine nucleotide dissociation inhibitors (GDIs), GEFs maintain a homeostatic balance of Rho-GTPase activity[4]. Each DOCK protein subclass contains additional functional domains conferring tissue specific expression and functional roles.

Class A and B DOCKs have established roles in neuromuscular health and function. DOCK1 and DOCK5 are critically important for myoblast fusion in the developing zebrafish [5]. A *DOCK1* variant has been correlated with age at loss of ambulation in DMD [6]. Case studies of human patients with *DOCK3* mutations highlight hypotonia and ataxia among other symptoms [7]. Additionally, our lab has previously shown that DOCK3 is involved in skeletal muscle metabolism by trafficking GLUT4 to the cell membrane and is also a dosage-sensitive modifier of Duchenne muscular dystrophy (DMD) pathology [8, 9]. While Class C DOCKs, DOCK6 and DOCK8, are primarily expressed in immune cells [10, 11], DOCK7 is highly expressed in skeletal muscle and neurons [12]. Most reported *DOCK7* pathogenic variants in ClinVar present with brain malformations and epileptic encephalopathy, 10% also display hypotonia or ataxia, and are diagnosed as developmental and epileptic encephalopathy-23 (DEE23)[13, 14]. Case studies of patients with *DOCK7* pathogenic variants also report, hypotonia and gross and fine motor delays [15–17]. Of all the DOCK family members, DOCK7 has the unique ability to activate both RAC1 and CDC42. By looping the DHR2 domain it achieves dual substrate specificity [18]. This dual activation potential leads to many critical roles in neuronal development such as directing axon polarity, promoting migration of interneurons and cortical neurogenesis based on the GEF’s ability to function as a sensitive molecular switch [19–21].

Within the skeletal muscle, proper regulation of Rho-GTPase activity by DOCK proteins is critical to maintain muscle function. RAC1 is necessary for both embryonic and fetal myogenesis [22, 23]. RAC1 activity is highest during fusion events likely due to its role in SCAR and WASP dependent actin polymerization [24]. RAC1 also signals through the dystrophin-glycoprotein complex (DGC) to promote c-JUN phosphorylation and cell growth [25]. RAC1 activity also controls AChR clustering at the post-synaptic neuromuscular junction both due to downstream activation of PAK1 and modulation of WNT signaling factors [26, 27]. DOCK proteins interact with RAC1 at the cellular membrane, and dystrophin-deficient myofiber membranes are significantly compromised thereby impairing RAC1 activation. We have previously shown that DOCK3 is a driver of DMD disease pathogenesis and that haploinsufficiency of *Dock3* improved outcomes in *mdx^5cv^* mice[28]. Similarly, expression analysis has revealed that DOCK proteins are increased in DMD patient muscle biopsies *and Dock7 in the mdx^5cv^* mouse and [29, 30]. Subsequently, the underlying disease mechanism of DOCK7 function in nerve and skeletal muscle has not yet been studied.

Here, we investigated the role of DOCK7 function in nerve and muscle through the generation and characterization of two conditional *Dock7* knockout mouse models. *Dock7* skeletal myofiber (*Dock7* mKO; *HSA*-Cre) and motor neuron (*Dock7* vKO; *vChAT*-Cre) were generated to define the roles of DOCK7 in each lineage. Muscle functional assessments were performed on the *Dock7* mKO and *Dock7* vKO mice to assess muscle performance and physiological strength. Additional evaluation of RAC1 activation and RNA-sequencing in each *Dock7* conditional mouse KO revealed a dysregulation of the WNT signaling pathway and other muscle structural transcripts. Haploinsufficiency of *Dock7* was performed by mating each conditional *Dock7* KO line with the *mdx^5cv^* (dystrophin-deficient) model and subsequently evaluated the resulting *Dock7* mKO:*mdx^5cv^* and *Dock7* vKO:*mdx^5cv^* KO. These double knockout (dKO) mice demonstrated an improvement of the overall DMD disease. Our studies advance the functional role of DOCK7 in normal muscle function and establish what pathways potentially disrupted by DOCK7 dysregulation in DMD.

## Results

### *Dock7* transcript is elevated in DMD patient and mouse skeletal muscles

Given our previous work identifying DOCK factors as secondary drivers of neuromuscular disorders, we evaluated the expression of *Dock7* transcript in *mdx* (*DBA/2J* strain) and *mdx^5cv^* (*C57BL/6J* strain) mouse skeletal muscles. In the tibialis anterior (TA) muscles of both dystrophin-deficient strains, we observed a greater than 1.5-fold increase in expression when normalized to their respective WT strain controls (**Figure 1A**). In the scMuscle single cell sequencing mouse muscle database[31], we observed elevated *Dock7* expression in muscle stem cells during activation through early fusion (**Figure 1B)**. These findings align with our hypothesis that DOCK7 (and other *DOCK* factors) are increased in expression during muscle injury, damage, or disease state. DOCK7 is highly expressed in both the skeletal myofibers and motor neurons, so to dissect the tissue-specific roles we utilized a conditional *Dock7* knockout mouse (*Dock7*^flox/flox^) that has loxP sites flanking exons 3 and 4 of mouse *Dock7[32]*. Previous work in *Dock7* mutants *Misty* and *Moonlight* demonstrated that coat color and bone density were impacted by loss-of-function variants [33]. In separate matings, we generated Dock7 skeletal myofiber (*Dock7* mKO) and motor neuron (*Dock7* vKO) knockout mice by mating the *Dock7*^flox/flox^ conditional mice to *HSA-*Cre (constitutive) and motor neuron *vChAT*-Cre mice to generate *Dock7* knockout mice in each respective lineage (**Figure 1C**). To evaluate *Dock7* expression, we performed digital droplet PCR (ddPCR) on cDNA prepared from the TA muscle. There is a significant reduction in *Dock7* transcription in both models with a greater reduction in the *Dock7* mKO muscle due to the presence of multiple cell types (**Figure 1D**). To confirm *Dock7* ablation, we performed PCR on genomic DNA extracted from mouse gastrocnemius, whole brain, and lumbar spinal cord tissues. We detected Cre-mediated recombination for the deleted *Dock7* exons 3 and 4 in the gastrocnemius of the *Dock7* mKO mice and in the lumbar spinal cord of *Dock7* vKO mice (**Figure 1E**). Some intact *Dock7* non-recombined gDNA could be detected in the *Dock7* vKO spinal cord which can be attributed to the presence of sensory neurons and interneurons. The *Dock7* mKO and *Dock7* vKO mice showed no visible phenotypes upon initial inspection and were similar in weights throughout their lifespan (**Data Not Shown**).

**Figure 1.**
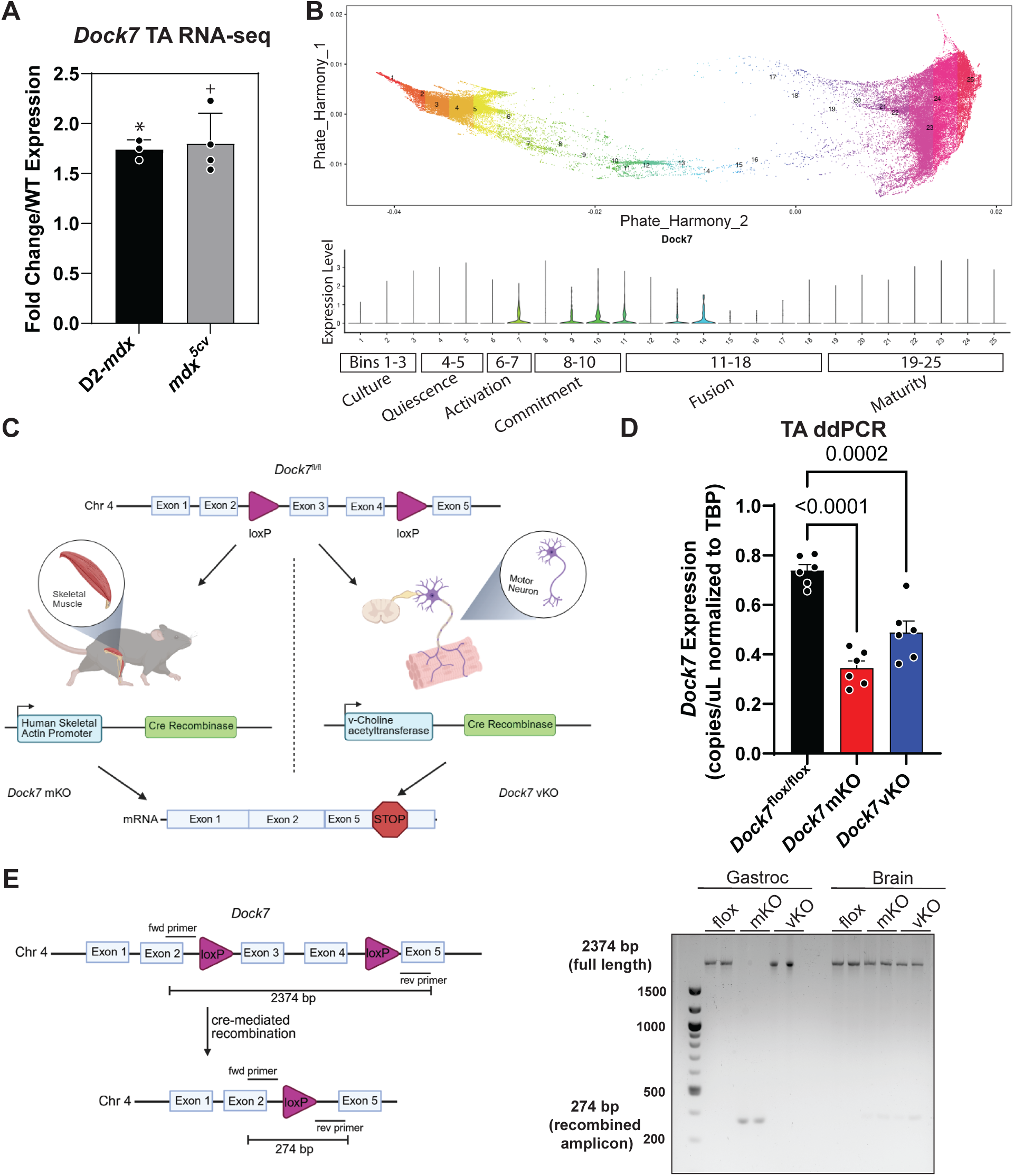
Generation and characterization of *Dock7* myofiber and motor neuron knockout mice. **A.** RNA sequencing from 6-month-old male D2-*mdx* and *mdx^5cv^* mice compared to their strain controls shows 1.5-2-fold upregulation of Dock7 transcript. TA muscle reveals *Dock7* expression is upregulated in two mouse models of DMD *q = 2.8E-04, +q = 1.82E-07. **B.** Transcriptional MuSC states identified in single cell sequencing of cultured MuSCs accessed from the scMuscle database displays upregulated *Dock7* expression primarily during activation through early fusion. Primary mouse muscle satellite cells (MuSCs) were cultured and sequenced to develop a detailed database of transitional in myoblast differentiation, then classified numerically and by color reflecting the stage of differentiation. Cell populations are organized into bins withtransitional states indicated by number and color. **C.** Schematic detailing the *Dock7* mKO and *Dock7* vKO models using the Cre-lox system to remove exons 3-4 and generate a premature stop codon in *Dock7* exon 5 **D.** *Dock7* mRNA expression in *Dock7*^flox/flox^, *Dock7* mKO, and *Dock7* vKO models normalized to *Tbp* (housekeeping control) levels. **E.** PCR primers were designed surrounding the recombination locus; full length intact DNA returns a product of 2374 bp and recombined DNA product is 274 bp in length. Genomic DNA from *Dock7* mKO gastrocnemius and *Dock7* vKO spinal cord show recombination with brain tissue as a control.

### Histological analysis reveals disruption of overall fiber size and RAC1 activation after *Dock7* ablation

We next evaluated the histopathological features of *Dock7* ablation in TA muscles from adult myofiber and motor neuron knockout mice. The *Dock7* mKO mice showed minimal pathologies with only small pockets of smaller, atrophied myofibers. Interestingly, the Dock7 vKO mice show larger myofibers throughout the TA muscles compared to Cre negative *Dock7*^flox/flox^ mice (denervation causes atrophy) (**Figure 2A**). Cross-sectional analysis (CSA) of the three cohorts showed that both the *Dock7* mKO and *Dock7* vKO mice had an overall shift in the total number of smaller myofibers compared to control mice (**Figure 2B**). Minimum Feret’s diameter also showed a trend toward a lower overall number of smaller myofibers (**Figure 2C**). RAC1 is an essential regulator of skeletal muscle growth, metabolism, and overall function[34, 35]. DOCK7 has been shown to be an essential regulator of RAC1 activation in multiple tissue types through an essential binding to its DOCK homology region 2 (DHR-2) domain[36]. We evaluated the levels of activated (RAC1) GTP within the TA muscles of the *Dock7* mKO, *Dock7* vKO, and *Dock7*^flox/flox^ control mice (**Figure 2D**). Quantification of RAC1 activity using a magnetic bead-based pulldown assay from each of the 3 cohorts showed reduced levels of active GTP-RAC1 in the *Dock7* mKO and *Dock7* vKO mice (**Figure 2E**).

**Figure 2.**
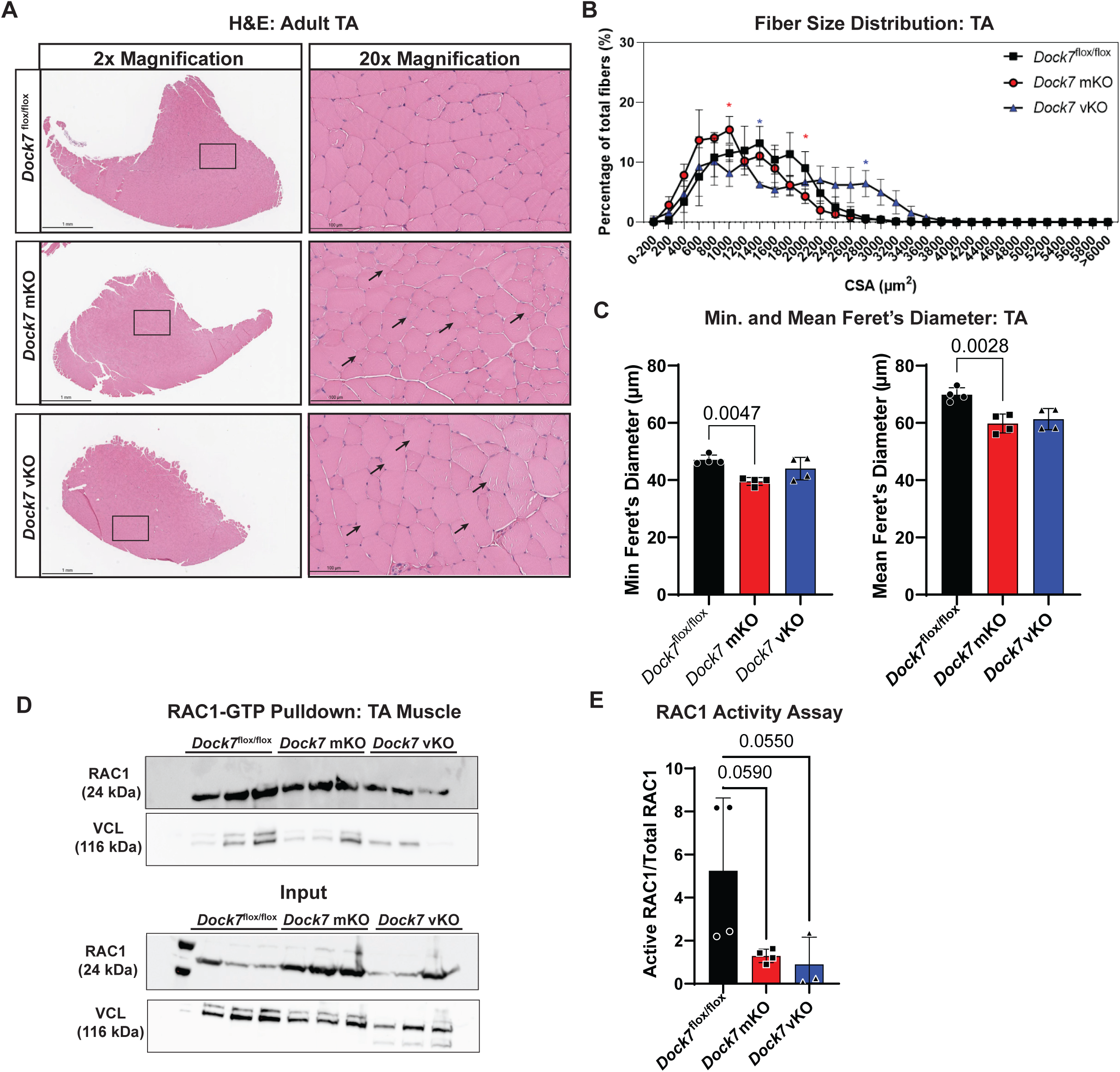
*Dock7* ablation in myofibers and motor neurons impairs muscle growth and RAC1 activation. **A.** Adult (4-month-old) H&E staining of *Dock7*^flox/flox^, *Dock7* mKO, and *Dock7* vKO TA muscles shows smaller, atrophied and denervated myofibers (black arrowheads) in *Dock7* mKO and *Dock7* vKO mice at 2x and 20x magnification. Scale bars = 1 mm and 100 µm. **B.** Fiber size CSA distribution was quantified using ImageJ and significance was determined using multiple t-tests corrected with false discovery rate, asterisks represent significance against *Dock7*^flox/flox^ mice. **C.** Minimum and mean Feret’s diameter was also measured in ImageJ, significance was determined using a one-way ANOVA with Bonferroni post-hoc analysis. **D.** GTP-RAC1 expression via a RAC1 pull-down assay in TA muscle lysates from the 3 cohorts. Expression and normalized to vinculin (VCL) control and total input. **E.** RAC1 activity assay quantified by western immunoblot densitometry. Significance determined using one-way ANOVA with Bonferroni post hoc *p<0.05, **p<0.01, ***p<0.001, ****p<0.0001.

### Conditional *Dock7* myofiber and motor neuron ablation leads to motor function deficits

Previous work identified DOCK7 as a mediator of neuronal axon guidance through its activation of RAC1 and cellular polarity[37]. We postulated that the genetic loss of *Dock7* in skeletal myofibers and/or motor neurons would have a significant impact on locomotor and muscle function. Forelimb grip strength measurements in adult mice revealed a significant drop in total normalized grip strength force in the *Dock7* mKO and *Dock7* vKO mice compared to the control cohort (**Figure 3A**). Open field test (OFT) of basal locomotion revealed decreased total distance traveled and average velocity in the *Dock7* mKO and *Dock7* vKO mice compared to the *Cre* negative *Dock7*^flox/flox^ mice (**Figure 3B**). We evaluated gait patterns using an automated Catwalk platform and observed gait defects in both the *Dock7* mKO and *Dock7* vKO mice (**Figure 3C**). Gait analysis of paw statistics, step sequence, and support (left and right, front, and hind paws) was conducted using Noldus Catwalk XT 10.6 (**Figure 3D**). *Dock7* vKO mice displayed significant disruptions to their gait pattern primarily presenting as decreased usage of the right hind paw and compensatory increase in use of the left front paw and significant time spent standing on ipsilateral paws with many additional disruptions (**Supplemental Figure 1**). *Dock7* mKO mice displayed a slight increase in usage of the left front paw (**Figure 3D**).

**Figure 3.**
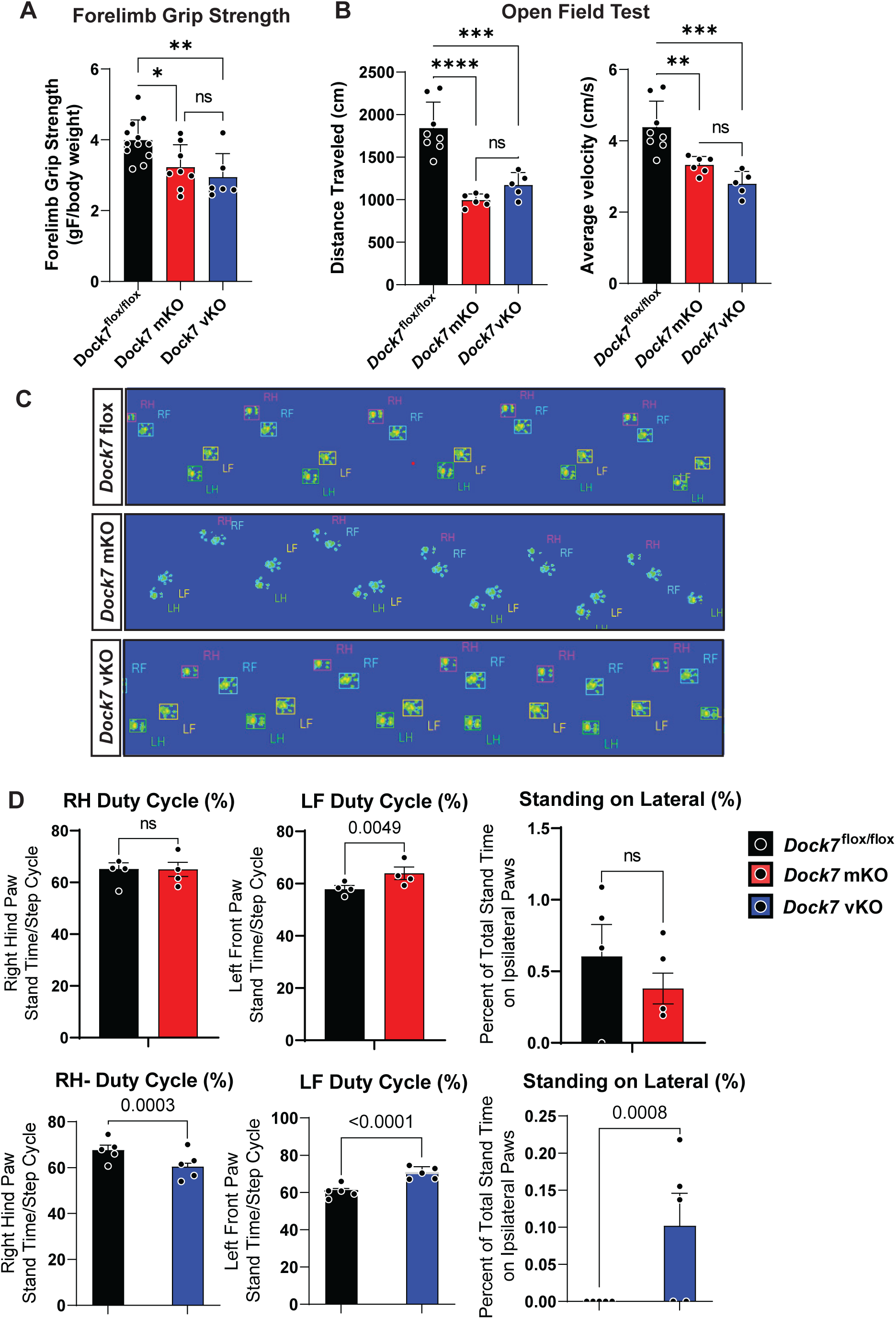
*Dock7* conditional knockout mice have muscle weakness, locomotor deficits, and gait disruption. **A.** Forelimb grip strength (grams Force/mouse body weight) measurements in the adult *Dock7*^flox/flox^ (black), *Dock7* mKO (red), and *Dock7* vKO (blue) mice. **B.** Open field-testing showing distance traveled (cm) and average velocity (cm/s) in each of the 3 cohorts. **C.** Representative catwalk gait traces showing a wider, disrupted gait in the *Dock7* vKO mice and no differences between the *Dock7* mKO and *Dock7*^flox/flox^ (control) cohorts. **D.** Many catwalk gait characteristics were measured between the 3 cohorts. Decreased right hind paw (RH) and increased left front paw (LF) duty cycle percentage as well as significant time spent standing on lateral paws indicate significant gait disruptions in the *Dock7* vKO mice but not the *dock7* mKO mice compared to *Dock7*^flox/flox^ (control). Significance determined using one-way ANOVA with Bonferroni post hoc *p<0.05, **p<0.01, ***p<0.001, ****p<0.0001. N = 6-10 mice per cohort.

### RNA-seq of TA muscles reveals significant disruption of WNT signaling, metabolic, and cytokine related transcripts

To identify the signaling pathways disrupted by *Dock7* loss, we performed RNA-sequencing on the TA muscles of adult *Dock7* mKO and *Dock7* vKO mice. We identified 488 differentially expressed genes (DEGs) in the *Dock7* mKO muscles compared to Cre-negative controls. Conversely, despite having a more severe gait disruption, only 120 DEGs were identified from the *Dock7* vKO mice (**Figure 4A**). Between the two cohorts were 20 DEGs in common in both *Dock7* conditional KO TA muscles (**Supplemental Table 1**). Volcano plots highlighting the DEGs observed in the *Dock7* mKO TA revealed differential expression of key metabolic and cell signaling transcripts, whereas the *Dock7* vKO TA volcano plot showed differential expression of metabolic and cytokine transcripts (**Figure 4B**). A heat map analysis showed the top 20-differentially regulated transcripts in *Dock7* mKO and vKO mouse muscles (**Figure 4C**). Ingenuity Pathway Analysis (IPA) of the *Dock7* mKO mice showed microfilament, cell adhesion, and myofibril assembly as the top biological processes disrupted (**Figure 4D**). IPA analysis of the *Dock7* vKO dysregulated pathways showed plasminogen, long chain fatty acid cellular import, and chemotaxis pathways as disrupted (**Figure 4E**). Together these analyses reveal key cell signaling, chemotaxis, and cellular migration pathways were disrupted by the absence of *Dock7* expression within the myofibers and motor neurons.

**Figure 4.**
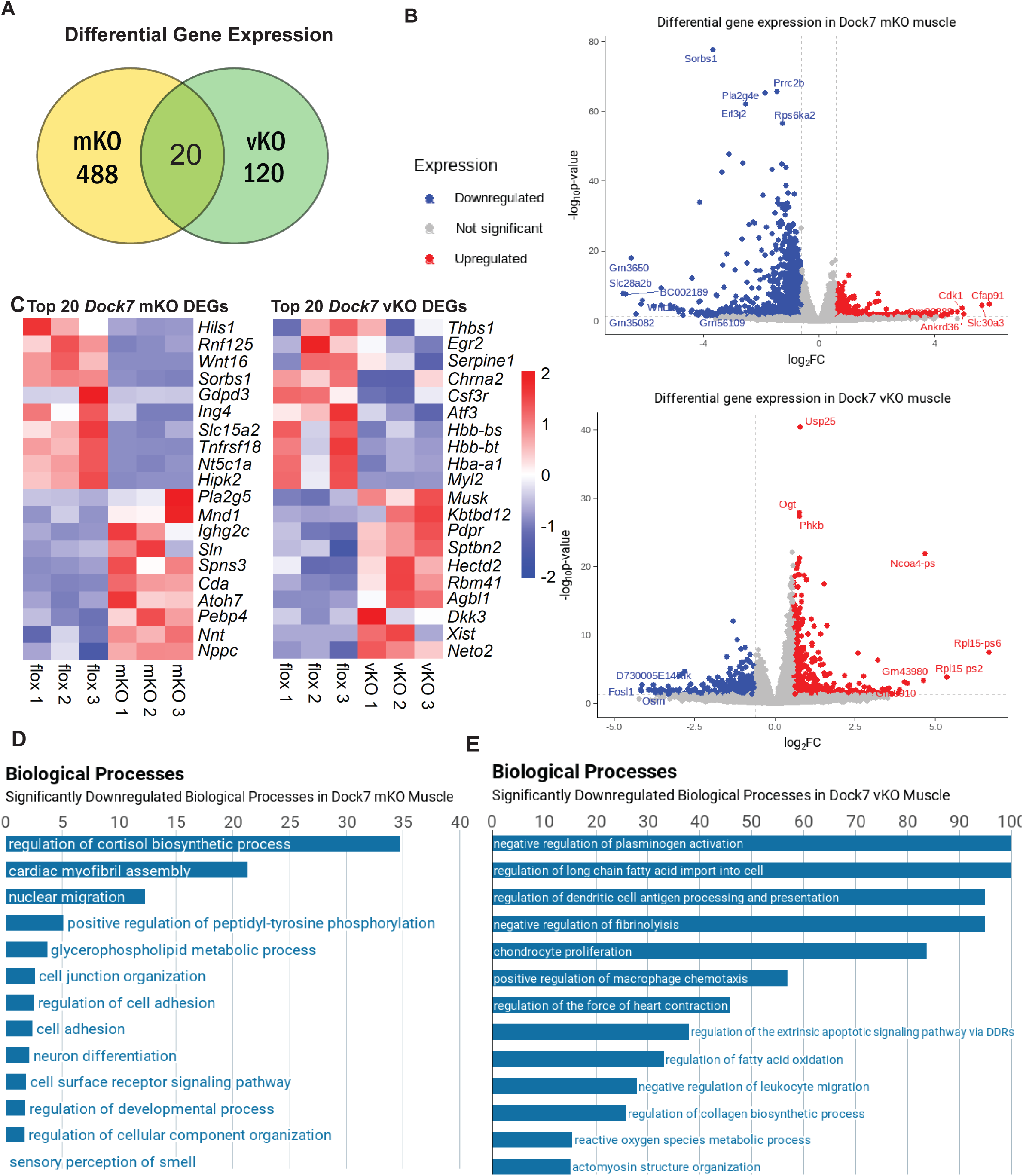
*Dock7* muscle and motor neuron ablation disrupts WNT signaling, cell migration, and cellular signaling pathway expression. **A.** Venn diagram shows the number of differentially expressed transcripts in the TA muscles in both *Dock7* cohorts. **B.** Volcano plots highlighting significantly downregulated (blue) and upregulated (red) genes in the TA muscles of 2 tissue specific knockout models *Dock7* mKO, and *Dock7* vKO. **C.** Heatmaps showing the top 20 most significantly up and downregulated differentially expressed genes (DEGs) in the TA muscles of *Dock7* mKO and *Dock7* vKO mice compared to the *Dock7*^flox/flox^ control cohort. **D-E.** Biological processes highlighting the top gene ontology (GO) pathways that were dysregulated in the muscles of the *Dock7* mKO and *Dock7* vKO mice.

### Disruption of WNT signaling is accompanied by significant disruption of motor junctions

We further probed the disruption of WNT signaling transcripts in the *Dock7* mKO and *Dock7* vKO mice as our transcriptomic analysis showed significant disruption in this pathway known to regulate muscle regeneration and axon guidance. Heat map analysis of 18 known WNT signaling transcripts that were dysregulated in either the *Dock7* mKO, *Dock7* vKO mice or both models. This revealed a significant upregulation of both canonical and non-canonical WNT transcripts in the *Dock7* vKO muscle and downregulation of WNT transcripts in *Dock7* mKO muscle (**Figure 5A**). Further examination of 5 WNT transcripts known to regulate skeletal muscle regeneration, MuSC proliferation, and DMD pathogenesis (*Lrp4*, *Wnt5a*, *Wnt2b*, *Wnt16*, and *Gsk3b*) were shown to be differentially expressed between the *Dock7* mKO and *Dock7* vKO gastrocnemius and soleus muscles using qPCR (**Figures 5B and 5C**). WNT pathway analysis revealed a significant disruption in key WNT upstream molecules in the *Dock7* mKO muscles, and more downstream WNT pathway regulation disruption in the *Dock7* vKO muscles (**Supplemental Table 2**). We evaluated the neuromuscular junction (NMJ) formation in *Dock7* mKO and *Dock7* vKO mouse muscles using immunofluorescent markers. Bungarotoxin and SH3/SV2A immunofluorescent labeling of the NMJ’s from the *Dock7* mKO and *Dock7* vKO mouse EDL muscles revealed a decrease in total area and increased fragmentation in the *Dock7* mKO mice and less of a significant decrease in NMJ clustering in the *Dock7* vKO EDL (**Figures 5C and 5D**). There was less of a significant decrease in NMJ clustering in the *Dock7* vKO mouse muscles compared to the Cre-negative cohorts (**Figures 5D**). These results suggest that the defect in the *Dock7* mKO mice may be attributed partially to malformed NMJ structures, whereas the motor neuron *Dock7* vKO mice may have muscle functional defects not related to NMJ formation.

**Figure 5.**
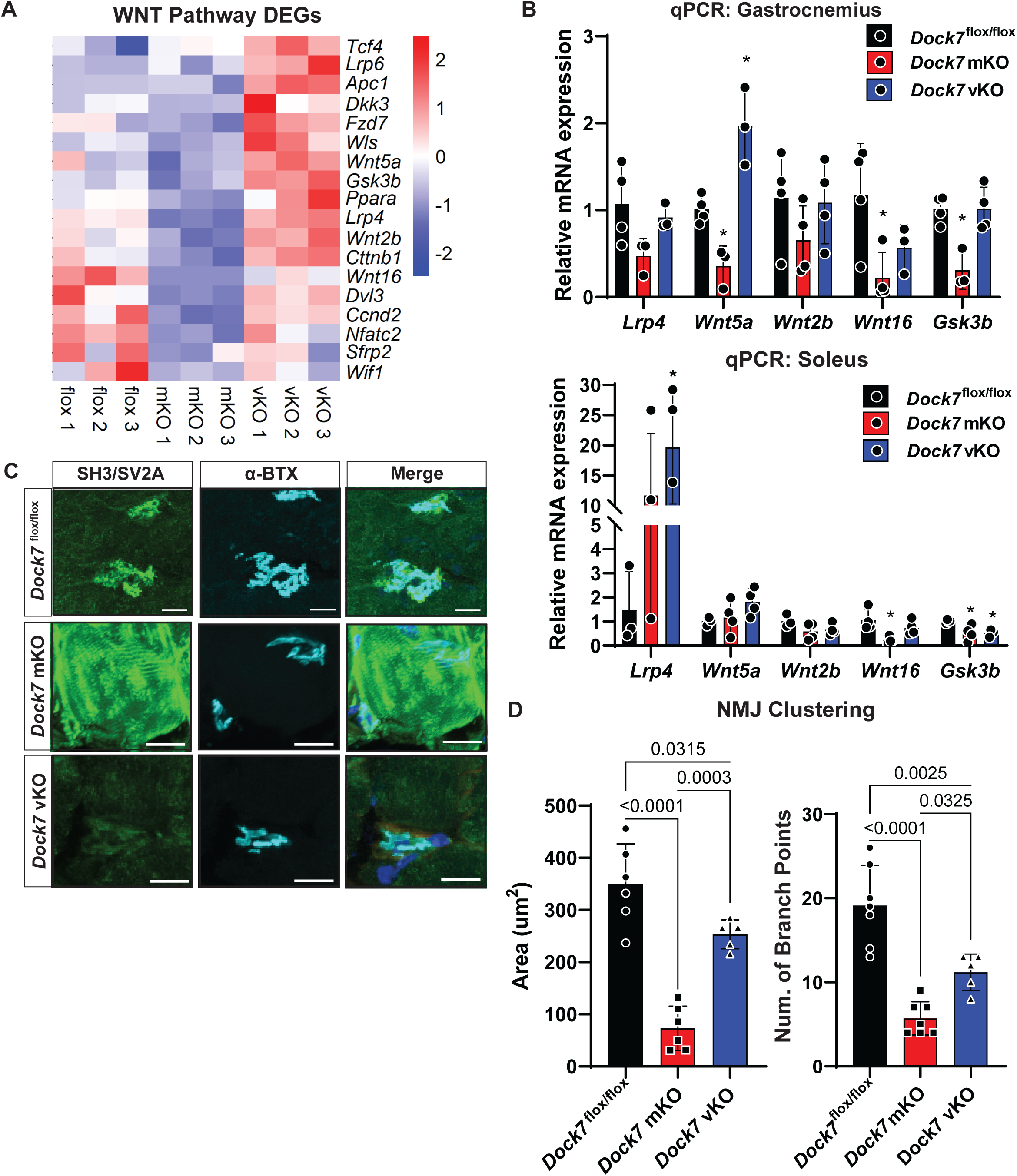
WNT pathway expression and activity is decreased in the skeletal muscles of *Dock7* mKO mice. **A.** Heatmap expression of WNT pathway intermediates showing decreased expression in *Dock7* mKO muscle and increased expression in *Dock7* vKO muscles. **B.** Quantitative PCR (qPCR) of *Lrp4*, *Wnt5a*, *Wnt2b*, *Wnt16*, and *Gsk3b* mRNA expression in the gastrocnemius and soleus muscles of the *Dock7*^flox/flox^, *Dock7* mKO, and *Dock7* vKO mice. Relative expression levels normalized to the *Actb* loading control. **C.** Immunofluorescence images of the EDL NMJs stained with SH3/SV2A, α-Bungarotoxin (BTX), and the merged images. Scale bars at 10 µm (*Dock7*^flox/flox^) and 20 µm (*Dock7* mKO and *Dock7* vKO). **D.** Quantification of NMJ clustering showing % area (µm^2^) and number of branch points quantified from z-stack images. Note the significant decrease in size of AChR clusters and the number of branch points. N = 5 mice/cohort.

### *Dock7* haploinsufficiency models counteract DMD symptoms and improves *mdx^5cv^* pathology

Given the upregulation of DOCK7 in DMD, we mated the *mdx^5cv^* mice to the *Dock7* mKO and *Dock7* vKO and evaluated their skeletal muscle histopathologies. Histopathological analysis of the adult TA muscles from the *Dock7* mKO: *mdx^5cv^* and *Dock7* vKO: *mdx^5cv^* showed no significant exacerbation of the *mdx^5cv^* muscle necrosis, but smaller overall myofibers with increased cellular infiltration indicative of elevated inflammation (**Figure 6A**) No changes were observed in the percentage of centralized myonuclei or percent cross-sectional area between the *mdx^5cv^*, *Dock7*: *mdx^5cv^* mKO, and *Dock7* vKO: *mdx^5cv^* mice (**Figures 6B and 6C**). Both minimum and mean Feret’s diameter showed no differences between the *mdx^5cv^* mice and the *Dock7* mKO: *mdx^5cv^*, and *Dock7* vKO: *mdx^5cv^* mice (**Figure 6B**). No changes were observed in the percentage of centralized myonuclei or percent cross-sectional area between the *mdx^5cv^*, *Dock7* mKO: *mdx^5cv^*, and *Dock7* vKO: *mdx^5cv^* mice (**Figures 6C and 6D**). Forelimb grip strength showed no difference between the *mdx^5cv^, Dock7* mKO: *mdx^5cv^*, and *Dock7* vKO: *mdx^5cv^* mice as did open field basal locomotion assessments (**Figures 6E and 6F**). Together, these show that dual loss of *Dock7* (both muscle and motor neuron) in dystrophin-deficient mice has no exacerbating impact on dystrophic symptoms. Our previous work demonstrated that *Dock3* haploinsufficiency improved DMD pathologies and outcomes in dystrophic mice[28]. Remarkably, both the *Dock7*^fl/+^ mKO: *mdx^5cv^* and *Dock7*^fl/+^ vKO: *mdx^5cv^* mice showed a substantial reduction in dystrophic pathological features compared to *mdx^5cv^* controls (**Figure 7A**). Minimum and mean Feret’s diameter was increased in the *Dock7*^fl/+^ mKO: *mdx^5cv^* and *Dock7*^fl/+^ vKO: *mdx^5cv^* cohorts compared to both *mdx^5cv^* and *Dock7*^flox/flox^ controls (**Figure 7B**). Additionally, we observed a shift towards larger myofiber sizes via CSA measurements and a reduction in the total number of centralized myonuclei (**Figures 7C and 7D**). Forelimb grip strength and open field testing showed improved muscle force and locomotor activity in *Dock7*^fl/+^ mKO: *mdx^5cv^* and *Dock7*^fl/+^ vKO: *mdx^5cv^* cohorts compared to both *mdx^5cv^* and *Dock7*^flox/flox^ controls (**Figures 7E and 7F**). Collectively, these studies reveal that haploinsufficiency of *Dock7* expression in dystrophin-deficient myofibers or motor neurons improves DMD-associated symptoms.

**Figure 6.**
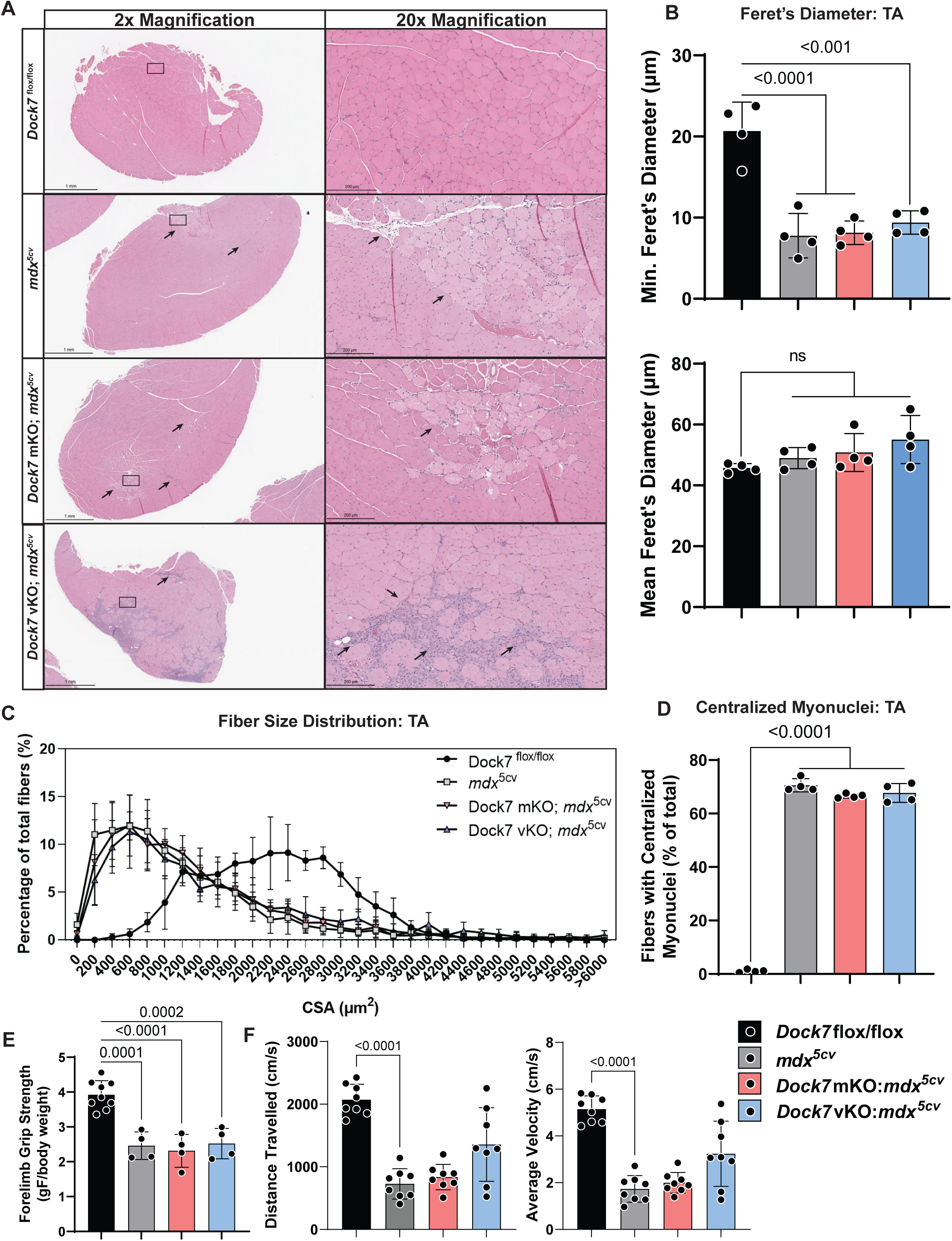
*Dock7* ablation does not exacerbate dystrophic skeletal muscle pathologies. **A.** H & E staining of adult 6-month-old mouse TA muscles revealed no significant differences in histopathology of the *Dock7* mKO; *mdx^5cv^* or *Dock7* vKO; *mdx^5cv^* double knockout mice. The *Dock7*^flox/flox^ control mice are used as controls. Scale bars = 1 mm and 200 µm. Arrows indicate areas of necrosis and cellular infiltration. **B.-D.** Centralized myonuclei, fiber size CSA, and minimum/mean Feret’s diameter between the 4 cohorts showed no exacerbation of the *mdx^5cv^* (DMD) muscle fiber pathologies. **E.-G.** Forelimb grip strength (grams Force/mouse body weight), distance traveled (cm), and average velocity (cm/s) in each of the 4 cohorts. Significance determined using one-way ANOVA with Bonferroni post hoc *p<0.05, **p<0.01, ***p<0.001, ****p<0.0001. N = 8-12 mice/cohort.

**Figure 7.**
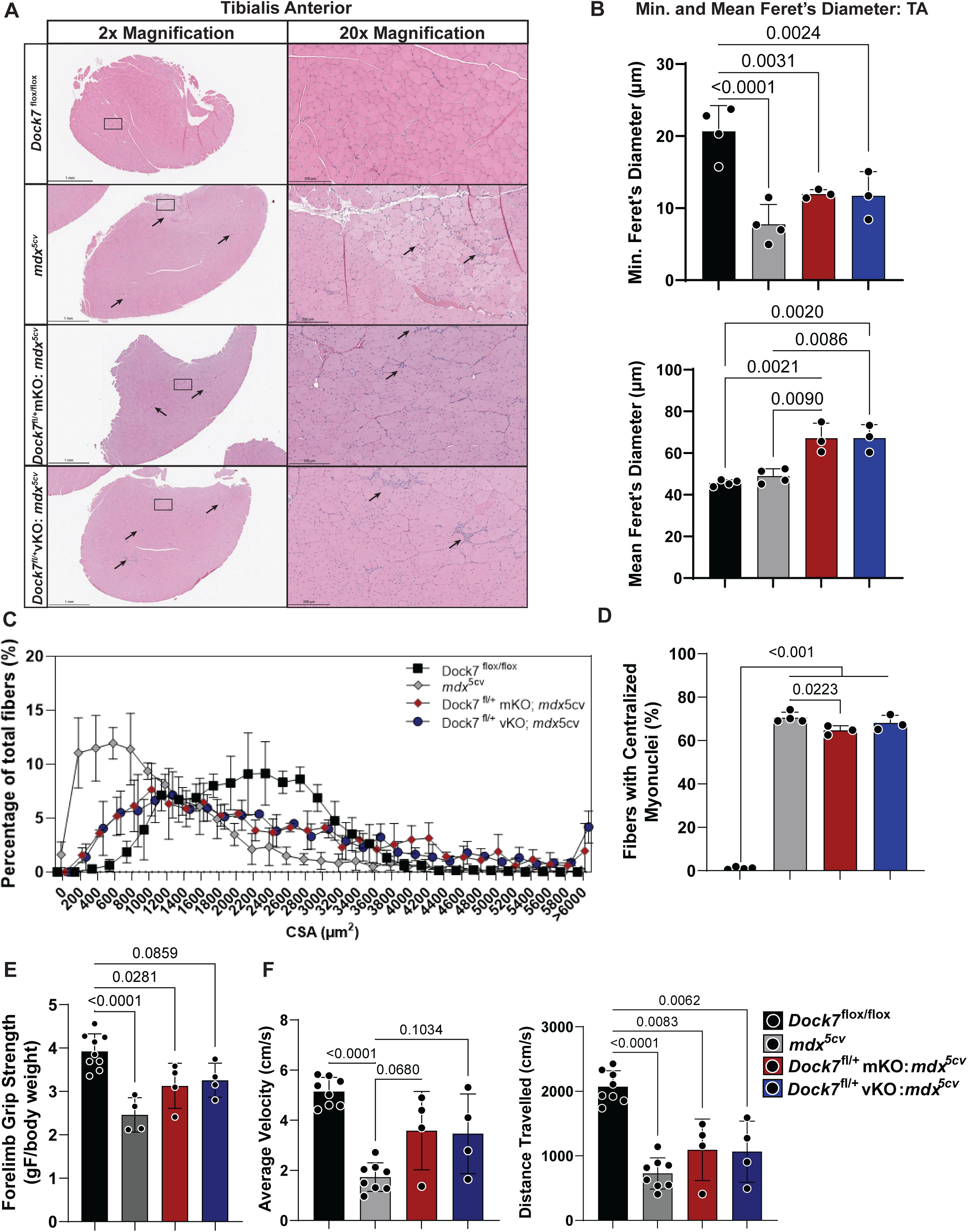
*Dock7* haploinsufficiency significantly improves DMD skeletal muscle pathologies. **A.** H&E histochemistry of adult 4-month-old TA muscles from *Dock7*^flox/flox^, *mdx^5cv^*, *Dock7*^fl/+^ mKO: *mdx^5cv^*, and *Dock7*^fl/+^ vKO: *mdx^5cv^* mice. Magnification shown at 2x and 20x insets with areas of necrotic tissue highlighted by arrowheads (black). Scale bars = 1 mm and 200 µm. **B. and C.** Minimum and mean Feret’s diameter and CSA shows *Dock7* haploinsufficiency in muscle with significant increases in Feret’s diameter and significant decreases in the number of small myofibers compared to *mdx^5cv^* mice N = 4 mice/cohort analyzed in each of the 4 cohorts: *Dock7*^flox/flox^ (black), *mdx^5cv^* (grey), *Dock7*^flox/+^ mKO:*mdx^5cv^* (red), and *Dock7*^flox/+^ vKO:*mdx^5cv^* (blue) mice. **D.** Centralized myonuclei counted as percentage of TA myofibers shown in each of the 4 cohorts. **E. and F.** Forelimb grip strength (grams Force/mouse body weight), distance traveled (cm), and average velocity (cm/s) in each of the 4 cohorts. Significance determined using one-way ANOVA with Bonferroni post hoc *p<0.05, **p<0.01, ***p<0.001, ****p<0.0001. N = 4-8 mice/cohort.

## Discussion

Our studies reveal that DOCK7 is an important regulator of skeletal muscle growth and development and impacts overall muscle function. The conditional *Dock7* myofiber KO mice had a slightly more severe pathological features than the conditional *Dock7* motor neuron KO mice. This suggests that DOCK7 may have a more substantial role in the skeletal muscle sustaining muscle function and development/growth and regulating post-synaptic NMJ function/formation. Global *Dock7*-deficiency in mice results in impaired bone and metabolic health, both symptoms that are associated with DMD. Our results indicate that DOCK7 muscle function may be impaired by defects in both RAC1 activation and WNT pathway signaling. RAC1 activation is essential for normal skeletal muscle growth, metabolism, and function and while there are several activators of RAC1, DOCK7 may play a unique role in RAC1 activation in muscle. Interestingly, while complete *Dock7* ablation in the myofiber and motor neuron had no impact on dystrophic symptoms, *Dock7* haploinsufficiency improved outcomes. These results implicate the dosage-sensitive requirement for DOCK7 expression within the muscle and offer a mechanism by which DMD symptom mitigation could be achieved via modulation of DOCK7 expression. One can envision gene therapy using AAV CRISPRi or PROTAC-mediated DOCK7 reduction as a novel therapeutic approach for DMD.

Questions remain with regard to the overlapping, and tissue-specific functions of the DOCK proteins in muscle and other tissues. This study showed the *Dock7* mKO mice had more of a disrupted NMJ structure and clustering, whereas the *Dock7* vKO mice had less of an impact on the NMJ and a greater impact on muscle function. DOCK7 protein shows high expression in neuronal tissues but also has broad expression in mesenchymal and other tissues. Conditional deletion and/or overexpression of *Dock7* may yield clues to its contribution to disease pathogenesis and secondary signaling modulation. DOCK7 expression in bone and fat tissues have been used as biomarkers for metabolic dysfunction, and similar diseases in which DOCK7 is disrupted may provide insight into upstream and downstream modulation of DOCK7 towards restoring its expression to homeostatic levels.

## Supporting information

All supplemental files

## Author Contributions

K.G.E., A.E.T-M, and M.S.A. conceived and planned the experiments. K.G.E., S.N.R., M.A.L., and M.K. completed all data collection and analysis. K.A.B. provided the conditional *Dock7*^flox/flox^ mice and consulted on results. K.G.E., A.E.T-M. and M.S.A. edited and wrote the manuscript. All authors discussed results and approved the manuscript prior to submission.

## Acknowledgments and Funding

We would like to thank members of the Alexander lab and UAB Center for Exercise Medicine for their critical evaluation of our manuscript. We would also like to thank members of the Alexander and Lopez labs for reviewing our manuscript and assistance with data interpretation. M.S.A. is supported by NIH grants from the Office of the Director under award number (U54OD030167) and an R21 grant from National Institute of Neurological Disorders and Stroke (NINDS) (R21NS140497). M.S.A. is supported by a Congressionally Directed Medical Research Program (CDMRP) Department of Defense (DoD) grant (MD240021). M.A.L. is supported by an NIH NIAMS grant (K08NS120812). K.A.B. is funded by a NIH National Institute of General Medical Sciences (NIGMS) Centers of Biomedical Research Excellence (COBRE) grant (P20GM152330). A.E.T-M. is supported by a NIH National Institute on Aging (NIA) R01 grant (R01AG075059). K.G.E. is supported by an NIH Eugene Kennedy Shriver National Institute of Child Health and Human Development (NICHD) training grant (T32HD071866).

## Methods

### Mice

WT (*C57BL/6J* strain; stock# 000664), *HSA*-Cre79 (B6.Cg-Tg(*ACTA1*-Cre)79Jme/J (Jax Mice stock# 006149), and vChAT (*ChAT*-IRES-Cre; Jax Mice stock# 006410) male and female mice were originally purchased from Jackson Labs (Bar Harbor, ME, USA). Conditional *Dock7* mice were obtained from Kathleen Becker and have loxP sites flanking mouse *Dock7* exons 3 and 4 and have been previously described[32]. All mice were maintained at UAB under standard housing and feeding conditions under an approved protocol (IACUC #21393). Mice were genotyped using site-specific primers and genomic DNA was isolated using Viagen Lysis Buffer (Viagen Biotech Inc.; Los Angeles, CA; Cat# 301-C). PCR genotyping was performed using Phusion High-Fidelity Polymerase (New England Biolabs, M0530S) along with primers Dock7GT forward 5’-GCCCAGATCACCCTTGTCTC-3’ and Dock7GT reverse 5’-GCA GCCAAGAGGGCAAAA-3’ flanking the floxed deletion region. The full-length WT allele was detected at 2374 bp and the recombination product was detected at 274 bp. All mice were housed under sterile, pathogen-free conditions with ad libitum access to food and water.

### Motor Assessments

All locomotor assessments were conducted in accordance with TREAT-NMD standard operating protocols for dystrophic mice[38]. Open field tests for basal locomotion were conducted using Noldus Ethovision version 17 (Noldus; Leesburg, VA). Mouse forelimb grip strength was conducted with Chatillon digital force gauge (Ametek STC; Edgewood, NY). Rotarod (San Diego Instruments; San Diego, CA) was conducted with on mice accelerating from 4 to 40 rpm over a period of 4 min after 3 consecutive days of training at 4 rpm. Gait analysis was performed on a Catwalk XT (Noldus), raw data was processed using the Catwalk XT 10.6 software (Noldus; Leesburg, VA).

### Histological Analysis

Skeletal muscle histological samples were fixed in 10% formalin, cut at 4 µm thickness, and stained for H&E or trichrome as previously described[39] using Histowiz histopathology services (Long Island City, NY). Some muscle biopsies were directly processed by slow freezing in Tissue-Tek O.C.T. (Sakura FineTek USA; Torrance, CA). Minimal Feret diameter, fiber size distribution, cross-sectional area, percent fibrotic area, and centralized nuclei were manually calculated as previously described using Fiji/ImageJ software[40, 41]. Neuromuscular junctions and other muscle sections were imaged using a Nikon AXR Confocal w/NSPARC (Leica Microsystems, Inc.; Buffalo Grove, IL) at 20X and 60X magnification. Briefly, extensor digitorum longus muscle was embedded in OCT (Fisher Scientific, Cat# 23-730-571; Hampton, NH) and sectioned longitudinally at 15 μm using cryostat (Leica CM1950; Leica Microsystems; Buffalo Grove, IL). Sections were post-fixed in 4% paraformaldehyde and dual-stained for Sv2A/2H3 (Cell Signaling Technology, Cat. #66724; Danvers, MA) and Developmental Studies Hybridoma Bank (DSHB) (Cat #531793; Iowa City, IA) and α–bungarotoxin (ThermoFisher Scientific, Cat. #B35451; Waltham, MA) for 24 hours after blocking with M.O.M buffer (Vector Laboratories, Cat. #MKB-2213-1; Newark, CA).

### Western Blotting

Protein lysates were isolated using RIPA Buffer (ThermoFisher Scientific, Cat#89900; ; Waltham, MA) with 1X HALT Protease Inhibitor Cocktail EDTA-free (ThermoFisher Scientific, #78425; Waltham, MA). Protein Lysates were quantified using a Pierce Bradford Colorimetric Assay Kit (ThermoFisher Scientific, Cat #23225; Waltham, MA). 50 μg of total protein lysate was loaded onto a 7.5% Mini-PROTEAN TGX Precast Protein gel (BioRad Laboratories; Cat# 4561023; Hercules, CA). Proteins were transferred to a 0.2 μm PVDF membrane (MilliporeSigma, Cat. #IPVH00010; Burlington, MA) and blocked in 5% milk in 0.1% TBS-T for one hour at room temperature. Then the membranes were incubated in primary antisera overnight at 4^0^C with gentle rocking. Membranes were washed in 0.1% TBS-T before being incubated with HRP-conjugated secondary antibodies (ThermoFisher Scientific; Cat# 31460; Waltham, MA) or Cell Signaling Technology, Cat# 7076; Danvers, MA) for 2 hours at room temperature with gentle rocking. After washing with TBS-T membranes were developed with ECL (Boston Bioproducts Inc.; Cat# WB-100-KIT; Milford, MA). Primary antibodies used include RAC1 (MilliporeSigma, Cat# 16-319; Burlington, MA), GSK3B (R&D Systems, Cat# MAB2506; Minneapolis, MN), WNT16 (ThermoFisher Scientific, Cat# MA5-47069; Waltham, MA), and Vinculin (Cell Signaling Technology; Cat# 4650; Danvers, MA).

### RAC1 Activity Assay

Samples were prepared using the RAC1 Activation Assay Kit (Millipore-Sigma, 17-283) according to kit instructions. Briefly, magnetic beads conjugated to the PAK1 protein binding domain were incubated with the sample. Western blot quantification was normalized to total RAC1 using anti-RAC1 clone 23A8 (MilliporeSigma; Cat# 16-319; Burlington, MA) levels of the input sample accounting for loading variability. Positive and negative controls were generated by saturating PAK1-PBD beads with the provided GDP and GTPγS control protein samples.

### RNA sequencing

Bulk RNA-sequencing of muscle samples were extracted using TRIzol (ThermoFisher Scientific; Cat#15596-026; Waltham, MA) and purified using Zymo Clean and Concentrate (Zymo Research Corporation; Cat# R1017; Irvine, CA) ensuring that all samples had a RIN score above 9, A260/280 >2 and A260/230 <2. Samples were submitted to Novogene (8801 Folsom Blvd #290, Sacramento, CA 95826) for bulk RNA sequencing with 30 million reads per sample. Data processing and differential gene expression were performed using Ingenuity Pathway Analysis (IPA). Secondary analysis of gene expression patterns were completed using the PANTHER Classification System[42]. All datasets were submitted to the NIH GeneExpressionOmnibus (GEO) under accession number GSE287712.

### Quantitative Real-Time PCR (RT-qPCR)

RNA samples were isolated using TRIzol (ThermoFisher Scientific; Cat#15596-026; Waltham, MA), quantified using a Nanodrop One UV-Vis Spectrophotometer (ThermoFisher Scientific; Cat# 13-400-518; Waltham, MA), then converted to cDNA using the Superscript IV First Strand Synthesis System (ThermoFisher Scientific; Cat# 18-090-010; Waltham, MA). Samples were added to a mastermix of Taqman probe sets (Wnt16: Mm00446420, Lrp4: Mm00554326, Wnt2b:Mm00437330, Wnt5a:Mm00437347, Gsk3b: Mm00444911 or Actb: Mm02619580) and 2x TaqMan Fast Advanced Master Mix (ThermoFisher Scientific, Cat# 4444557; Waltham, MA). Samples were run on a Roche LightCycler480 Real Time PCR (Roche Life Sciences, Cat# 05015243001; Basel, Switzerland) and results were quantified using the 2^−ΔΔCt^ method and normalized to β-actin as the loading control.

### Statistical Analysis

All statistical analysis was conducted in Graphpad Prism version 9 (Dotmatics; Boston, MA). Comparisons between 2 groups were performed with students t-test and Tukey post hoc test. Significance between 3 or more groups was determined with a two-way ANOVA and Bonferroni post hoc, *p<0.05, **p<0.01, ***p<0.001, ****p<0.0001.

## REFERENCES

1. Boland, A., J.F. Cote, and D. Barford, Structural biology of DOCK-family guanine nucleotide exchange factors. FEBS Letters, 2023. 597(6): p. 794.

2. Whitehead, I.P., et al., Dbl family proteins. Biochim Biophys Acta, 1997. 1332(1): p. F1–23.

3. Thompson, A.P., et al., RHO to the DOCK for GDP disembarking: Structural insights into the DOCK GTPase nucleotide exchange factors. J Biol Chem, 2021. 296: p. 100521.

4. Duman, J.G., et al., Mechanisms for spatiotemporal regulation of Rho-GTPase signaling at synapses. Neurosci Lett, 2015. 601: p. 4–10.

5. Moore, C.A., et al., A role for the Myoblast city homologues Dock1 and Dock5 and the adaptor proteins Crk and Crk-like in zebrafish myoblast fusion. Development, 2007. 134(17): p. 3145–53.

6. Weiss, R.B., et al., Long-range genomic regulators of THBS1 and LTBP4 modify disease severity in duchenne muscular dystrophy. Ann Neurol, 2018. 84(2): p. 234–245.

7. Iwata-Otsubo, A., et al., DOCK3-related neurodevelopmental syndrome: Biallelic intragenic deletion of DOCK3 in a boy with developmental delay and hypotonia. Am J Med Genet A, 2018. 176(1): p. 241–245.

8. Reid, A.L., et al., DOCK3 is a dosage-sensitive regulator of skeletal muscle and Duchenne muscular dystrophy-associated pathologies. Hum Mol Genet, 2020. 29(17): p. 2855–2871.

9. Samani, A., et al., DOCK3 regulates normal skeletal muscle regeneration and glucose metabolism. bioRxiv, 2023.

10. Xu, X., et al., LRCH1 interferes with DOCK8-Cdc42-induced T cell migration and ameliorates experimental autoimmune encephalomyelitis. J Exp Med, 2017. 214(1): p. 209–226.

11. Wilkie, H., et al., DOCK8 is essential for neutrophil mediated clearance of cutaneous S. aureus infection. Clin Immunol, 2023. 254: p. 109681.

12. Consortium, G.T., The Genotype-Tissue Expression (GTEx) project. Nat Genet, 2013. 45(6): p. 580–5.

13. Landrum MJ, L.J., Riley GR, Jang W, Rubinstein WS, Church DM, Maglott DR, ClinVar: public archive of relationships among sequence variation and human phenotype Nucleic Acids Research, 2014. 42(1): p. D980–5.

14. Samani, A., et al., DOCKopathies: A systematic review of the clinical pathologies associated with human DOCK pathogenic variants. Hum Mutat, 2022.

15. Bai, B., et al., Novel DOCK7 mutations in a Chinese patient with early infantile epileptic encephalopathy 23. Chin Med J (Engl), 2019. 132(5): p. 600–603.

16. Haberlandt, E., et al., Characteristic facial features and cortical blindness distinguish the DOCK7-related epileptic encephalopathy. Mol Genet Genomic Med, 2021. 9(3): p. e1607.

17. Perrault, I., et al., Mutations in DOCK7 in individuals with epileptic encephalopathy and cortical blindness. Am J Hum Genet, 2014. 94(6): p. 891–7.

18. Kukimoto-Niino, M., et al., Structural Basis for the Dual Substrate Specificity of DOCK7 Guanine Nucleotide Exchange Factor. Structure, 2019. 27(5): p. 741–748 e3.

19. Yang, Y.T., C.L. Wang, and L. Van Aelst, DOCK7 interacts with TACC3 to regulate interkinetic nuclear migration and cortical neurogenesis. Nat Neurosci, 2012. 15(9): p. 1201–10.

20. Watabe-Uchida, M., et al., The Rac activator DOCK7 regulates neuronal polarity through local phosphorylation of stathmin/Op18. Neuron, 2006. 51(6): p. 727–39.

21. Nakamuta, S., et al., Dual role for DOCK7 in tangential migration of interneuron precursors in the postnatal forebrain. Journal of Cell Biology, 2017. 216(12): p. 4313–4330.

22. Vasyutina, E., et al., The small G-proteins Rac1 and Cdc42 are essential for myoblast fusion in the mouse. Proc Natl Acad Sci U S A, 2009. 106(22): p. 8935–40.

23. Charrasse, S., et al., M-cadherin activates Rac1 GTPase through the Rho-GEF trio during myoblast fusion. Mol Biol Cell, 2007. 18(5): p. 1734–43.

24. Kaipa, B.R., et al., Dock mediates Scar- and WASp-dependent actin polymerization through interaction with cell adhesion molecules in founder cells and fusion-competent myoblasts. J Cell Sci, 2013. 126(Pt 1): p. 360–72.

25. Oak, S.A., Y.W. Zhou, and H.W. Jarrett, Skeletal muscle signaling pathway through the dystrophin glycoprotein complex and Rac1. J Biol Chem, 2003. 278(41): p. 39287–95.

26. Bai, Y., et al., Balanced Rac1 activity controls formation and maintenance of neuromuscular acetylcholine receptor clusters. J Cell Sci, 2018. 131(15).

27. Henriquez, J.P., et al., Wnt signaling promotes AChR aggregation at the neuromuscular synapse in collaboration with agrin. Proc Natl Acad Sci U S A, 2008. 105(48): p. 18812–7.

28. Reid, A.L., et al., DOCK3 is a dosage-sensitive regulator of skeletal muscle and Duchenne muscular dystrophy-associated pathologies. Human Molecular Genetics, 2020. 29(17): p. 2855–2871.

29. Lopez, M.A., et al., Smad8 Is Increased in Duchenne Muscular Dystrophy and Suppresses miR-1, miR-133a, and miR-133b. International Journal of Molecular Sciences, 2022. 23(14): p. 7515.

30. Sothers, H., et al., Late-Stage Skeletal Muscle Transcriptome in Duchenne Muscular Dystrophy Shows a BMP4-Induced Molecular Signature. Journal of Cachexia, Sarcopenia and Muscle, 2025. 16(4): p. e70005.

31. McKellar, D.W., et al., Large-scale integration of single-cell transcriptomic data captures transitional progenitor states in mouse skeletal muscle regeneration. Communications Biology, 2021. 4(1): p. 1280.

32. Bishop, K.A., et al., CRISPR/Cas9-Mediated Insertion of loxP Sites in the Mouse Dock7 Gene Provides an Effective Alternative to Use of Targeted Embryonic Stem Cells. G3 (Bethesda), 2016. 6(7): p. 2051–61.

33. Blasius, A.L., et al., Mice with mutations of Dock7 have generalized hypopigmentation and white-spotting but show normal neurological function. Proceedings of the National Academy of Sciences, 2009. 106(8): p. 2706–2711.

34. Sylow, L., et al., Rac1 Is a Novel Regulator of Contraction-Stimulated Glucose Uptake in Skeletal Muscle. Diabetes, 2013. 62(4): p. 1139–1151.

35. Raun, S.H., et al., Skeletal muscle Rac1 mediates exercise training adaptations towards muscle glycogen resynthesis and protein synthesis. Redox Biology, 2025. 86: p. 103844.

36. Kukimoto-Niino, M., et al., Structural Basis for the Dual Substrate Specificity of DOCK7 Guanine Nucleotide Exchange Factor. Structure, 2019. 27(5): p. 741–748.e3.

37. Watabe-Uchida, M., et al., The Rac Activator DOCK7 Regulates Neuronal Polarity through Local Phosphorylation of Stathmin/Op18. Neuron, 2006. 51(6): p. 727–739.

38. Willmann, R., et al., Enhancing translation: Guidelines for standard pre-clinical experiments in mdx mice. Neuromuscular Disorders, 2012. 22(1): p. 43–49.

39. Samani, A., et al., miR-486 is essential for muscle function and suppresses a dystrophic transcriptome. Life Science Alliance, 2022. 5(9): p. e202101215.

40. Schindelin, J., et al., Fiji: an open-source platform for biological-image analysis. Nature Methods, 2012. 9(7): p. 676–682.

41. Karuppasamy, M., et al., Conditional Dmd ablation in muscle and brain causes profound effects on muscle function and neurobehavior. Communications Biology, 2025. 8(1): p. 1736.

42. Mi, H., et al., Large-scale gene function analysis with the PANTHER classification system. Nature Protocols, 2013. 8(8): p. 1551–1566.

