## Supplementary material for "DOCK7 regulates WNT signaling and NMJ clustering in normal and DMD muscle": All supplemental files

**Supplemental Figure Legend**

**Supplemental Figure 1. Gait analysis reveals *Dock7* vKO mice have an abnormal gait and gait characteristics.** Several additional measures of gait were found to be significantly disrupted in the *Dock7* vKO mice using Catwalk digital gait analysis. Left front foot (LF) and right front foot (RF) characteristics are shown. Statistical significance was determined using a student's t-test with Tukey post hoc. N = 4 mice/cohort.

**Supplemental Table 1. Commonly dysregulated transcripts between *Dock7* mKO and *Dock7* vKO TA muscles.** The 20 commonly dysregulated transcripts identified between the *Dock7* mKO and the *Dock7* vKO TA muscles, fold change, and adjusted p-values are listed.

**Supplemental Table 2. Supplemental Table 2. WNT pathway transcripts disrupted in *Dock7* mKO and *Dock7* vKO mouse muscles.** Table depicting the differential expression of WNT pathway-related transcripts in *Dock7* mKO, *Dock7* vKO, and *mdx*$^{5cv}$ TA muscles. Fold change normalized to WT (C57Bl/6J) expression levels to determine if Up (>2.0 fold or greater), Down (-2.0 fold or greater), or no change in expression.

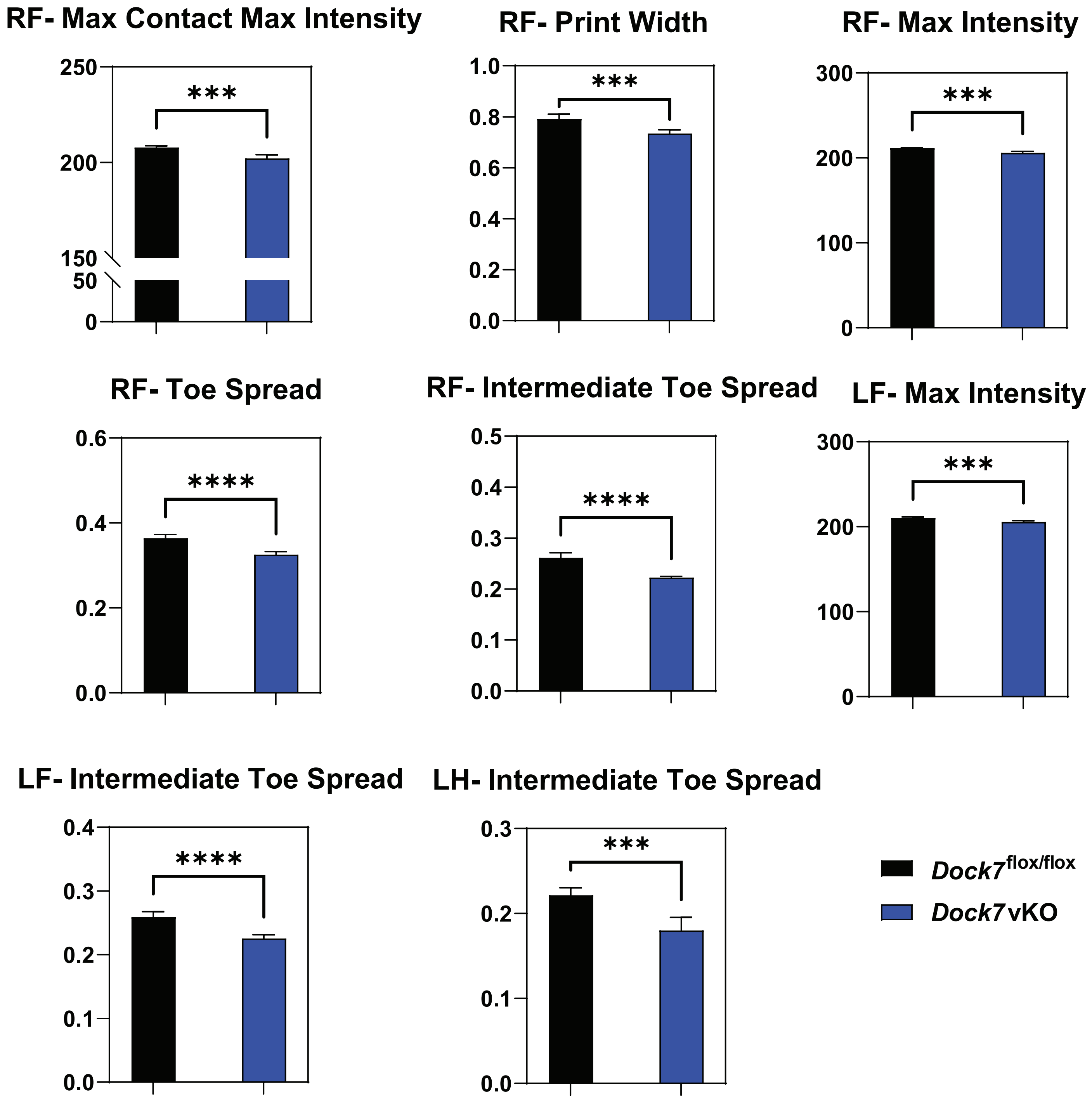


Supplemental Figure 1: Gait analysis reveals *Dock7* vKO mice have an abnormal gait and gait characteristics.

| Common Dysregulated Transcripts | | | | |
|---|---|---|---|---|
| | *Dock7* mKO | | *Dock7* vKO | |
| Gene Name | Fold Change | q value | Fold Change | q value |
| *Ighm* | 2.33505272 | 0.004147 | 2.01945481 | 0.018695 |
| *Neto2* | 2.12110299 | 0.006834 | 2.71509362 | 7.34E-05 |
| *Zfp750* | 0.48588633 | 0.00944 | 0.38380183 | 0.000189 |
| *AI838599* | 0.45844271 | 0.002461 | 0.43529309 | 0.000472 |
| *Csrp3* | 0.44320489 | 0.02423 | 0.4701468 | 0.036402 |
| *Rev1* | 0.41404126 | 9.92E-07 | 2.24818751 | 1.44E-07 |
| *Gm56916* | 0.39834543 | 0.006061 | 2.67951571 | 1.29E-05 |
| *Fan1* | 0.3408804 | 4.06E-10 | 2.06387012 | 8.67E-09 |
| *Dkk3* | 0.31159898 | 0.002867 | 2.4117473 | 0.027331 |
| *Gm8424* | 0.30200278 | 1.08E-11 | 2.35936468 | 8.35E-10 |
| *Adamts8* | 0.26942649 | 0.00037 | 0.39140965 | 0.011881 |
| *Pdpr* | 0.25620665 | 1.21E-18 | 2.90711534 | 2.26E-15 |
| *Myl2* | 0.23122477 | 0.043673 | 0.12693144 | 0.001863 |
| *Sned1* | 0.22315126 | 4.01E-13 | 2.01007956 | 3.87E-06 |
| *Gm4861* | 0.20639133 | 0.003119 | 3.30073678 | 0.000767 |
| *Gm44669* | 0.17121527 | 0.000801 | 2.24867213 | 0.025608 |
| *Thbs1* | 0.12771767 | 0.005409 | 0.16434574 | 0.014257 |
| *Pm20d2* | 0.11148318 | 1.06E-17 | 3.10439086 | 6.05E-10 |
| *Lrch3* | 0.1034146 | 1.02E-08 | 2.31057939 | 0.000957 |

**Supplemental Table 1: Commonly dysregulated transcripts between *Dock7* mKO and *Dock7* vKO TA muscles.**

| Gene Name | *Dock7* mKO | *Dock7* vKO | *mdx*$^{5cv}$ |
|---|---|---|---|
| Wnt2b | Down | Up | Down |
| Wnt5a | Down | Up | Down |
| Wls | Down | Up | Up |
| Wnt16 | Down | No Change | No Change |
| Dkk3 | Down | Up | Down |
| Sfrp2 | No Change | No Change | Up |
| Lrp4 | Down | Up | Down |
| Ccnd2 | Down | No Change | Down |
| Ppara | Down | Up | Down |
| Wif1 | Down | No Change | Down |
| Nfatc2 | Down | No Change | No Change |
| Lrp6 | No Change | Up | No Change |
| Tcf4 | No Change | Up | No Change |
| Fzd7 | Down | Up | No Change |
| Dvl3 | Down | Up | Up |
| Apc1 | No Change | Up | No Change |
| Cttnb1 | Down | Up | Down |
| Gsk3b | No Change | Up | No Change |

**Supplemental Table 2. WNT pathway transcripts disrupted in *Dock7* mKO and *Dock7* vKO mouse muscles.**
